# Analysis of small EV proteomes reveals unique functional protein networks regulated by VAP-A

**DOI:** 10.1101/2023.07.18.549588

**Authors:** Bahnisikha Barman, Marisol Ramirez, T. Renee Dawson, Qi Liu, Alissa M Weaver

## Abstract

Extracellular vesicles (EVs) influence cell phenotypes and functions via protein, nucleic acid and lipid cargoes. EVs are heterogeneous, due to diverse biogenesis mechanisms that remain poorly understood. Our previous study revealed that the endoplasmic rectiulum (ER) membrane contact site (MCS) linker protein VAP-A drives biogenesis of a subset of RNA-enriched EVs. Here, we examine the protein content of VAP-A-regulated EVs. Using label-free proteomics, we identified down- and up-regulated proteins in sEVs purified from VAP-A knockdown (KD) colon cancer cells. Gene set enrichment analysis (GSEA) of the data revealed protein classes that are differentially sorted to SEVs dependent on VAP-A. STRING protein-protein interaction network analysis of the RNA-binding protein (RBP) gene set identified several RNA functional machineries that are downregulated in VAP-A KD EVs, including ribosome, spliceosome, mRNA surveillance, and RNA transport proteins. We also observed downregulation of other functionally interacting protein networks, including cadherin-binding, unfolded protein binding, and ATP-dependent proteins.

**Significance of the study:** Little is known about biogenesis mechanisms that underlie EV heterogeneity. This study explores the protein repertoire of a specific subset of EVs that we recently identified to be generated at ER MCS and that are highly enriched in RNAs. We find that proteins from several classes of RNA machineries, including spliceosomes, are downregulated in EVs purified from cells knocked down for the ER MCS linker protein VAP-A. These data suggest that dynamic regulation of these protein machineries at ER MCS are involved in the sorting of RNA-RBP complexes into EVs.

## Introduction

EVs are secreted membrane-bound carriers that carry proteins, lipids, and nucleic acids to neighboring and distant cells to mediate cell-cell communication [1]. Small EVs are 30-150 nm in diameter and can be generated either as exosomes from endosomal sources or as ectosomes from the plasma membrane. The composition of small EVs is highly dependent on both cell type and biogenesis mechanism [1]). Recent data indicate that EVs released from cells are highly heterogeneous, with diverse functions dependent on their cargoes. We previously showed that the key ER MCS protein VAP-A, along with its binding partner CERT, drives the biogenesis of a specific RNA-containing class of small EVs [2]. Additionally, we found that the candidate RNA binding proteins (RBPs) Ago2, heterogenous nuclear ribonucleoprotein A2B1 (hnRNPA2B1) and synaptotagmin binding cytoplasmic RNA-interacting protein (SYNCRIP), are less abundant in VAP-A KD small and large EVs [2].

A recent study identified 1542 human RBPs [3-5] as involved in RNA processing, modification, and stability. RBPs also regulate nuclear RNA export, subcellular localization, and mRNA translation [3; 5]. One important function of EV-associated RBPs is to control the stability and sorting of the RNAs into the EVs [6]. We and others identified Argonaute 2 (Ago2) and other RISC complex proteins as potent mediators of miRNA sorting into EVs [7-11]. Other studies have shown that miRNAs with specific sequence motifs (i.e., GGAG and GGCU) are selectively sorted into EVs by hnRNPA2B1 and SYNCRIP [12; 13]. In addition, the RNA-binding protein Y-box I (YBX-1) has been shown to be involved in sorting of mRNAs, miRNAs, and other exRNAs into EVs [14-16]. While these studies highlight the importance of RBPs in RNA sorting into EVs, it is unclear which RNA-RBP complexes and machineries are involved in the sorting of RNA into EVs.

Here, we use a quantitative proteomic approach to identify significant differences in the protein cargos of small EVs derived from control and VAP-A KD DKs-8 colorectal cancer cells. We find that VAP-A-KD DKs8 SEVs are deprived of proteins associated with several functional categories, including protein folding chaperone, cadherin binding, ATP activity, and RBPs. Protein-protein interaction (PPI) network analysis of the RBP category by STRING revealed that downregulated RBPs in VAP-A KD DKs-8 small EVs are primarily associated with RNA transport, ribosomes, mRNA surveillance, and spliceosomes. Several identified downregulated proteins were further validated to be reduced in abundance in VAP-A KD EVs. These data are consistent with VAP-A controlling a subpopulation of small EVs originating at ER MCS and identify protein machineries that are preferentially sorted.

## Results and Discussion

To obtain highly pure small EVs, we subjected conditioned media collected from DKs-8 colon cancer cells to a series of centrifugations to remove debris and large EVs, followed by a cushion density gradient purification [17]. Consistent with our previous study, specific EV marker proteins segregated into density gradient fractions 6 and 7 (Figure 1A). Analysis of the combined fractions 6 and 7 confirmed a small but significant decrease in the number of small EVs purified from VAP-A KD cells by nanoparticle tracking analysis (Figure 1B and 1C, [2]). Transmission electron microscopy of negatively stained purified EV samples confirmed the purity and morphology of the EVs (Fig 1D).

**Figure 1.**
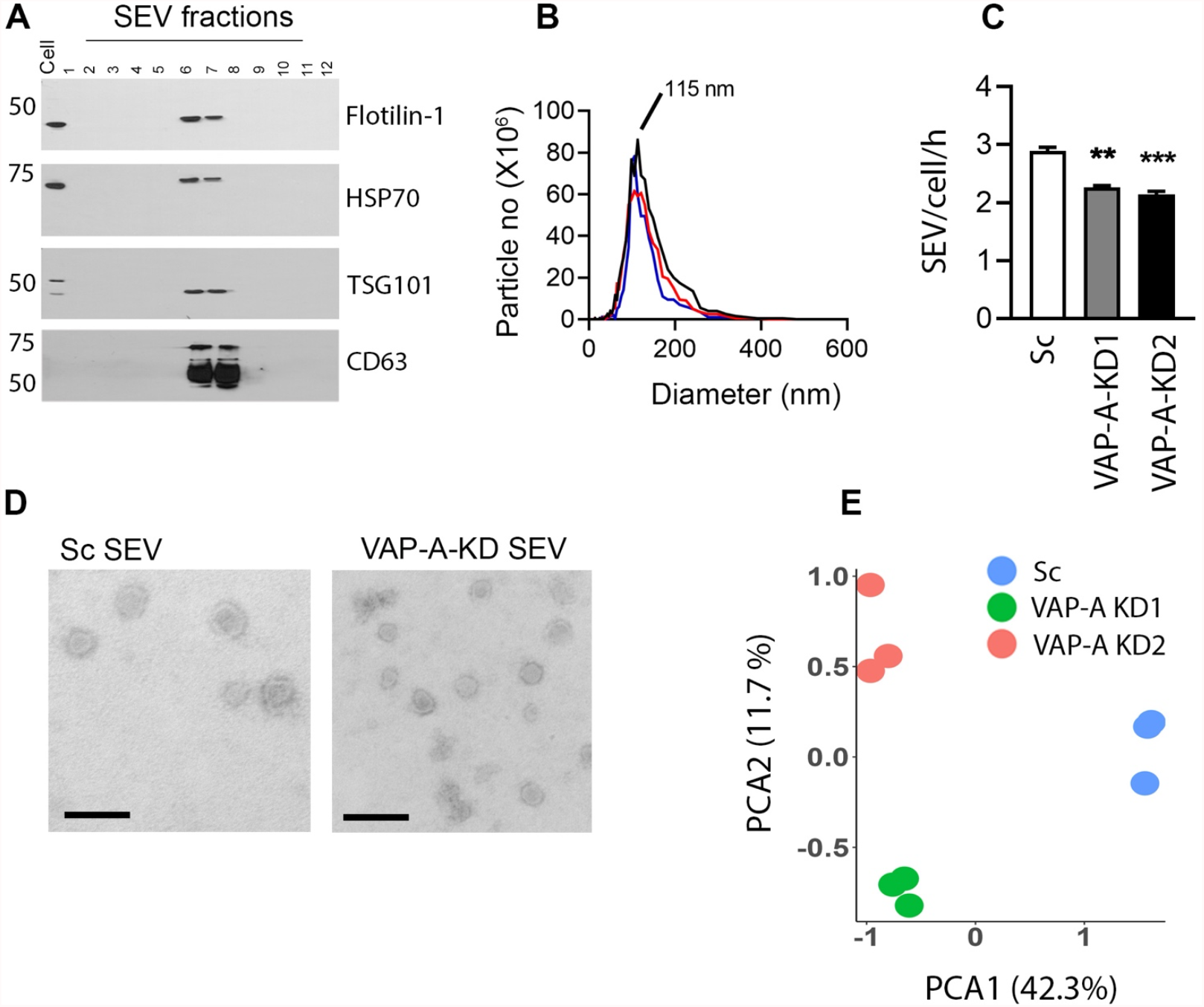
Characterizaon and proteomic analysis of VAP-A-KD small EVs in DKs-8 cells. A.Western blot analysis of control (Sc) DKs-8 cell lysates, and cushion density gradient fractions of small EVs for EV markers flollin-1, HSP70, TSG101, and CD63. B.Representave traces from nanoparticle tracking analysis of DKs-8 control (Sc) and VAP-A-KD (KD1 and KD2) SEVs (Median values from three independent experiments are ploted). C.Quantaon of vesicle secretion rate, calculated from nanoparticle tracking data for EVs collected in 48 hours from DKs8 cells (cell numbers quantitated at me of media harvest) (n=3). D. Representave TEM images for small EVs purified from DKs-8 control (Sc) and VAP-A-KD2 (KD) cells. Scale bar is 100 nm (n=3, dataset originally published in [2]) E. Principal component analysis (PCA) of protein composition in small EVs showing segregaon of KD (KD1 and KD2) from control (SC) data (n=3 biological replicates).

To identify the small EV protein cargos, equal amounts of protein extracted from control (Sc) and VAP-A-KD small EVs were subjected to label-free quantitative mass spectrometry (see Methods). Three independent experiments reliably identified over 1100 proteins (Supplemental Table 1). Principal component analysis (PCA) showed that the proteins are well segregated between control (Sc) and VAP-A-KD (KD1 and KD2) SEVs (Figure 1E). Using a criteria of the absolute log2-fold change (|log 2-fold change| > 1), we identified 225 proteins to be reduced and 129 proteins to be up-regulated in VAP-A-KD small EVs. One caveat is that many of these differences were not statistically significant due to low spectral counts for many of the proteins (Supplemental Table 1).

To identify enriched functions/pathways of sorted proteins in VAP-A-KD small EVs, we performed gene set enrichment analysis (GSEA). Several functions/pathways are significantly enriched in downregulated proteins in VAP-A KD small EVs (FDR<0.25), including adhesion, cell-cell junction, unfolded protein binding, and RNA binding (Table 1 and Supplemental Table 2). As we previously found that VAP-A controls biogenesis of a specific subpopulation of EVs that is enriched in RNA, we performed STRING analysis to identify PPI among the RNA binding proteins that are negatively enriched in VAP-A KD EVs according to the GSEA analysis. Several protein interaction clusters were identified, including ribosome, mRNA surveillance, RNA transport, and spliceosome pathways (Fig 2A and 2B). STRING PPI analysis on other downregulated classes of proteins, including unfolded protein, chaperone, cadherin binding, and ATP-dependent activity are shown in Supplemental Figure 1. The GSEA also identified functional classes that are enriched in upregulated proteins in VAP-A KD EVs, including extracellular matrix (ECM) proteins and inflammatory proteins (Table 2, Supplemental Table 3, and Supplemental Figure 2).

**Table 1:** Abbreviated list of GSEA pathways enriched in downregulated proteins in VAPA-KD small EVs. For each gene set, the pathway/function name, # of proteins, normalized enrichment score (NES), and false discovery rate (FDR) q value are listed.

| Function/Pathways | NES | # Number of Proteins | FDR q-value |
| --- | --- | --- | --- |
| CADHERIN_BINDING | -1.96 | 123 | 0.014 |
| CHROMATIN_BINDING | -1.8 | 40 | 0.058 |
| ESTABLISHMENT_OF_PROTEIN_LOCALIZATION_TO_ORGANELLE | -1.8 | 60 | 0.31 |
| CYTOKINESIS | -1.8 | 26 | 0.238 |
| MUSCLE_SYSTEM_PROCESS | -1.79 | 30 | 0.204 |
| ATP_DEPENDENT_ACTIVITY | -1.78 | 98 | 0.055 |
| IMPORT_INTO_NUCLEUS | -1.77 | 24 | 0.189 |
| UNFOLDED_PROTEIN_BINDING | -1.76 | 33 | 0.051 |
| CELL_CELL_JUNCTION | -1.76 | 83 | 0.101 |
| CELL_LEADING_EDGE | -1.75 | 86 | 0.081 |
| RNA_BINDING | -1.73 | 255 | 0.052 |
| EPIDERMIS_DEVELOPMENT | -1.72 | 40 | 0.249 |

**Figure 2.**
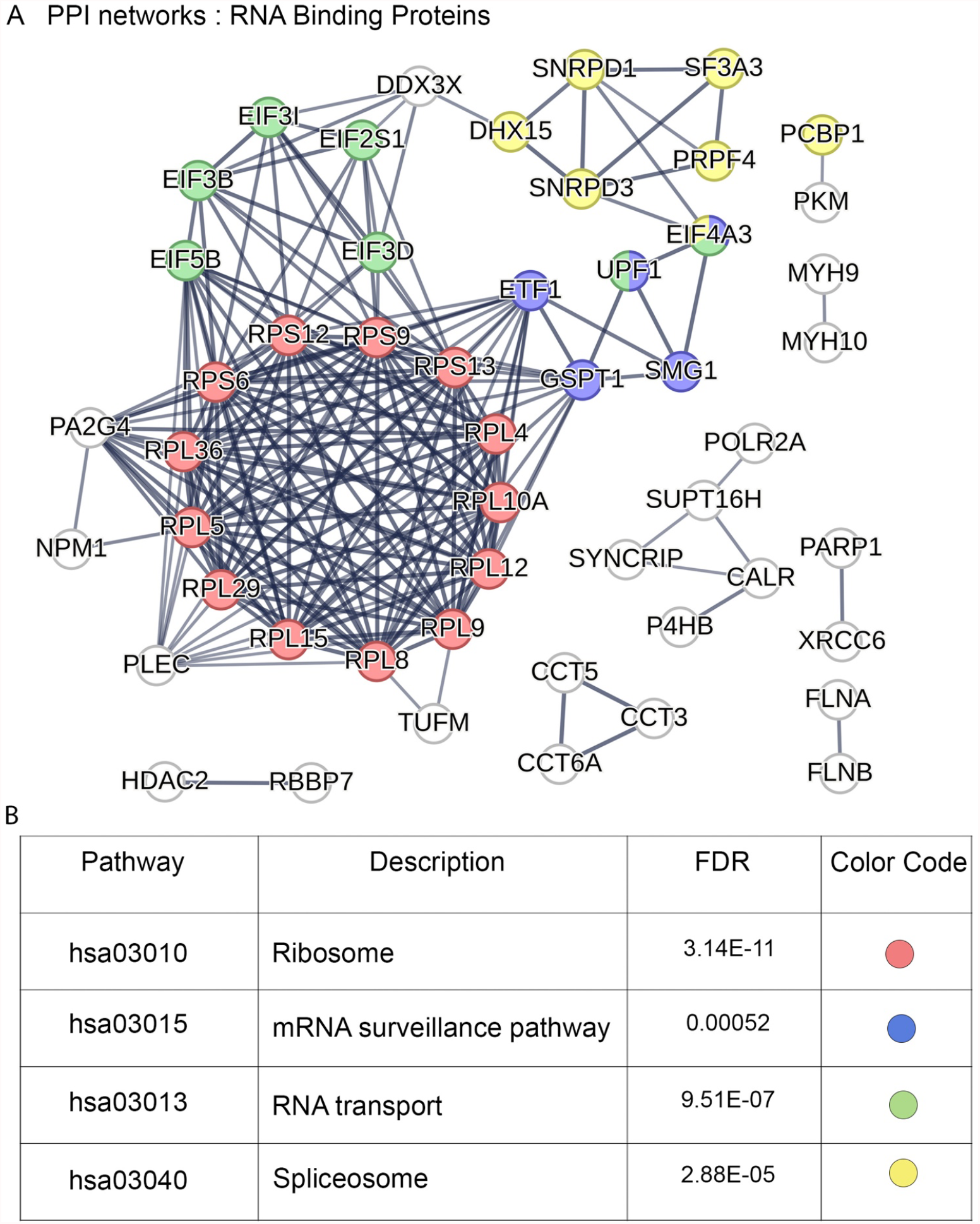
Idenfication of PPI RBP networks within the downregulated proteins for VAP-A-KD SEVs. A. STRING PPI Network of the GOMF_RNA binding proteins (GO:0003723) that were downregulated in VAP-A-KD SEVs. B.Table shows KEGG pathways and their FDR values. Color codes define different pathways (RED denotes Ribosome; BLUE denotes mRNA surveillance pathway; GREEN denotes RNA transport’; and YELLOW defines Spliceosome proteins).

**Table 2:**
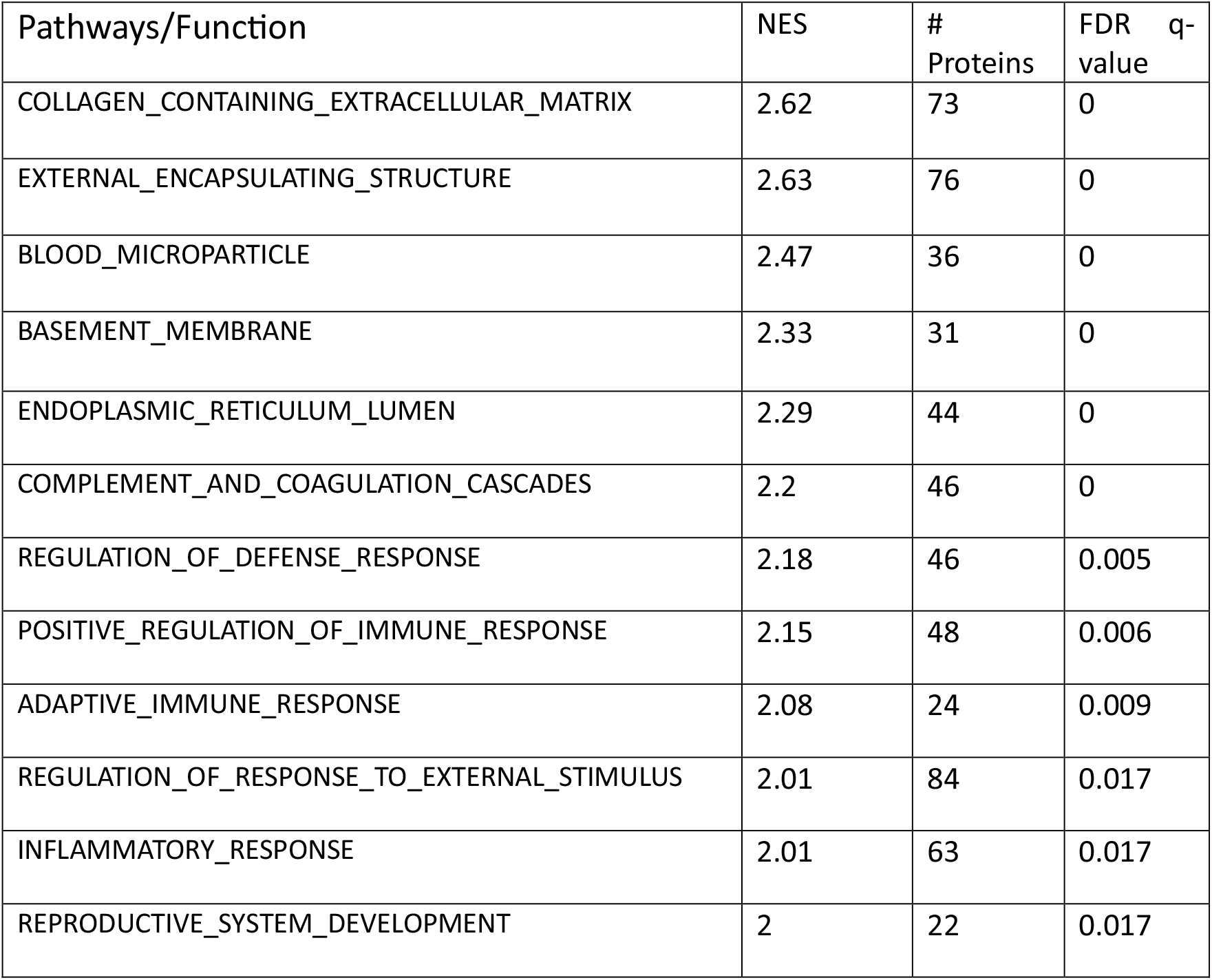
Abbreviated list of GSEA pathways enriched in upregulated proteins in VAPA-KD small EVs. For each gene set, the pathway/function name, # of proteins, normalized enrichment score (NES), and false discovery rate (FDR) q value are listed.

To validate a subset of the proteins identified as downregulated in VAP-A KD EVs, lysates of both small EVs and cells from control and VAP-A KD cells were analysed by Western blotting (Figures 3A and 3B). Two RBPs were selected for validation: the spliceosome component SF3A3 and EIF4A3, which is involved in multiple processes, including splicing regulation, mRNA surveillance, and RNA transport (Figure 2). Prominin-1 is a pentaspan cholesterol-binding protein that localizes to both the ER and plasma membrane and is often used as a stem cell marker [18; 19]. Western blot analyses validated that the abundance of these proteins is indeed lower in VAP-A KD small EVs (Figures 3A and 3B). There was either no change or upregulation of the same proteins in VAP-A KD cell lysates. Notably, EIF4A3 was upregulated in VAP-A KD cell lysates, indicating strong sorting to EVs. In addition, very little Prominin-1 was detected in cell lysates compared with EVs, consistent with it being strongly sorted to EVs [20].

**Figure 3.**
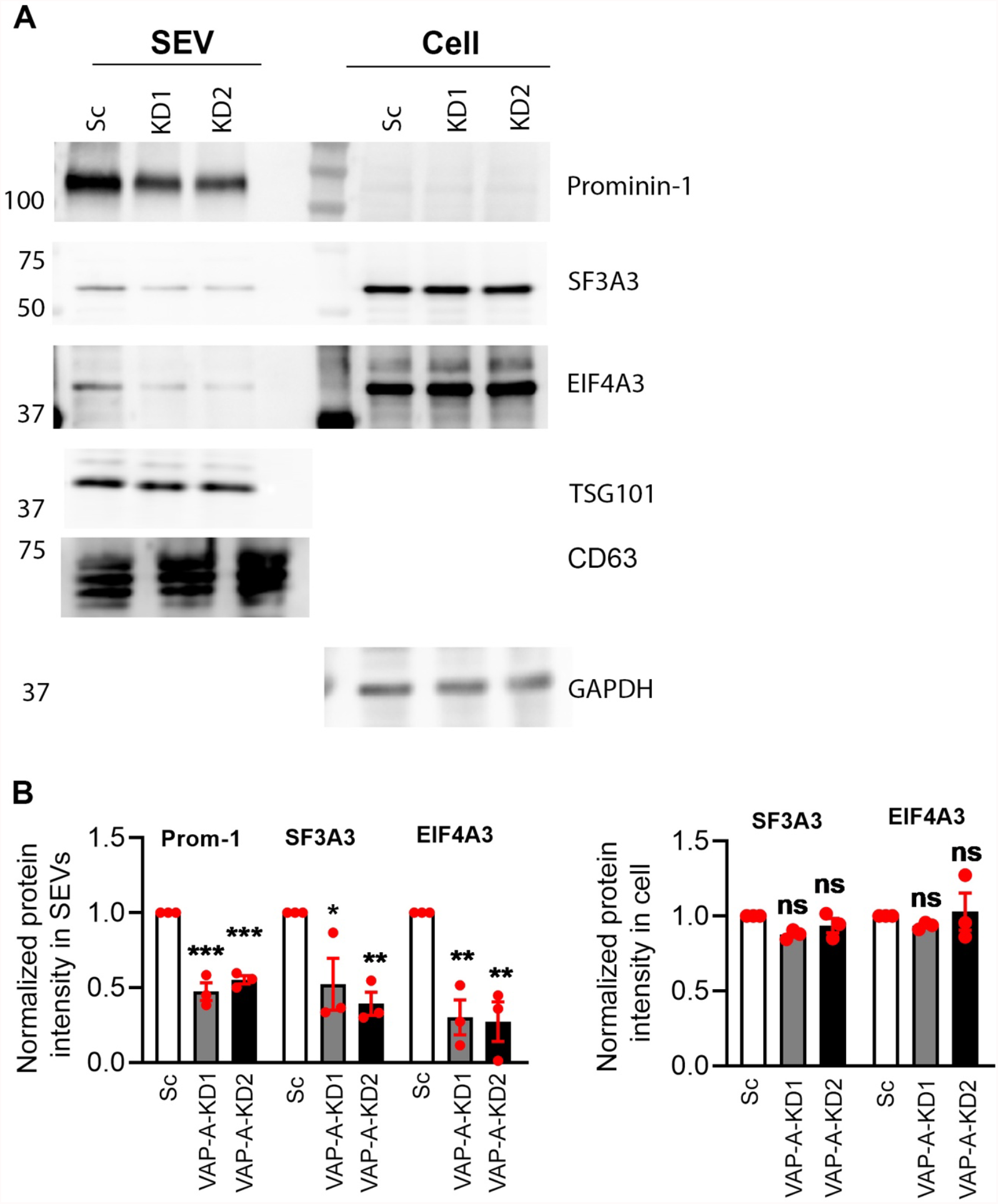
Western blot analysis of select downregulated proteins in control and VAP-A-KD SEVs and cells. A. Western blot analysis of SEVs purified from control (Sc) and VAP-A-KD (KD1 and KD2) DKs-8 cells and respective parental cells for RNA binding proteins SF3A3, EIF4A3, and Prominin-1. TSG101 was probed as a loading control for EVs, CD63 as an additional EV marker, and GAPDH as a loading control for cells. B. Quanfication of immunoblots from three independent experiments. Values are normalized by TSG101 for small EVs and by GAPDH for cell lysates. Data are plotted as mean ± SEM. ∗p < 0.05, ∗∗p < 0.01, ∗∗∗p < 0.001. ns, not significant.

Overall, these data are consistent with VAP-A controlling EV biogenesis-related processes at ER MCS, including cholesterol and ceramide transport and sorting of RNAs into EVs. The finding that spliceosome components are included in the downregulated RBPs in VAP-A KD cells is somewhat surprising, since these proteins primarily localize to the nucleus. Nonetheless, the nuclear spliceosome component SF3A3 is one of the strongest downregulated proteins in VAP-A KD EVs according to the proteomics analysis, a finding that we further validated by Western blot (Table 1 and Figure 3). SF3A3 forms the trimeric SF3A complex with SF3A1 and SF3A2 in the cytoplasm [21], which is then imported to the nucleus where it contributes to assembly of the active 17S U2 snRNP, a component of the major spliceosome complex [22]). While these functional complexes retain the vast majority of SF3A3 in the nucleus for splicing, a small amount of SF3A3 can be detected in the cytoplasmic fraction of cellular lysates [23] where it might be trafficked to sites of EV biogenesis. In fact, in the multiciliated cells of Xenopus epithelium, SF3A3 localizes to specialized cytoplasmic granules called dynein axonemal particles [24]. SF3A3 has also been shown to redistribute to the cytoplasm upon ubiquitination [23] and through interaction with the tumor suppressor protein CSR1 [25]. Lastly, it is possible that localization of spliceosome complexes to the ER/cytoplasm could be due to their reported involvement in nonsense mediated decay [26]. Indeed, our STRING analysis predicted that the downregulated spliceosome components may link to cytoplasmic mRNA surveillance through EIF4A3. Western blot analysis confirmed downregulation of EIF4A3 in VAP-A KD cells. EIF4A3 is an RNA helicase that shuttles between the nucleus and the cytoplasm and is part of the exon junction complex that controls mRNA splicing and monitors whether mRNA is appropriately spliced before translation [27]. EIF4A3 is also upregulated in several cancers [27].

In addition to RBPs, we found that the stem cell marker Prominin-1 is downregulated in VAP-A KD EVs. This might occur due to the cholesterol transfer activity of VAP-A at ER MCS [28; 29], as Prominin-1 is a cholesterol-binding protein [30-32]. Prominin-1 itself is known to induce biogenesis of small and large EVs [33], which may account in part for some of the decrease in EV numbers released from VAP-A KD cells (Figure 1, [2]). Cholesterol transfer activity may also relate to some of the upregulated proteins in VAP-A KD EVs. Notably, ApoE is a key lipoprotein constituent and binds to LRP [34], Supplemental Figure 3); their upregulation in VAP-A KD EVs may relate to cholesterol transport activity of VAP-A at ER MCS.

In summary, we identified classes of proteins, including RBPs, spliceosome, and mRNA surveillance machinery, whose presence in EVs is regulated by the ER MCS linker protein VAP-A. Future studies should investigate the mechanisms that govern their sorting into EVs.

## Experimental procedures

### Cell lines

DKs-8 colon cancer cells were cultured in DMEM (Corning) supplemented with 10% Fetal bovine serum (FBS), non-essential amino acids (Sigma), and L-glutamine. HEK293FT lentiviral packaging cells were cultured in DMEM supplemented with 10% FBS and 0.5 mg/ml G418 Sulfate (Corning). Stable shRNA scrambled control and shRNA VAP-A knockdown cell lines were previously described [2]. The shRNA constructs for VAP-A in pLKO.1 lentiviral shRNA expression system were purchased from Dharmacon (Cat #RHS3979-201759439 Cat #RHS3979-201759438). The scrambled control construct was acquired from Addgene (Plasmid no #26701).

### Extracellular vesicle isolation and nanoparticle tracking analysis

For the cushion density gradient method, cells were cultured at 80% confluence in serum-free DMEM. Aer 48 hours, the conditioned medium was collected from the cells and the EVs were isolated via serial centrifugation. Floating live cells, dead cell debris, and large EVs were respectively collected from the conditioned medium by centrifugation at 300 x g for 10 min, 2,000 x g for 25 min, and 10,000 x g (Ti45 rotor, Beckman Coulter) for 30 min. The supernatant was then overlaid onto a 2 ml 60% iodixanol cushion and spun at 100,000 x g (SW32 rotor, Beckman Coulter) for 18h. The bottom 3 ml, including the 1 ml of collected EVs + 2 ml iodixanol (40% iodixanol final concentration) were transferred to the bottom of another tube and then 20%, 10% and 5% iodixanol were layered successively on top. These iodixanol dilutions were prepared by diluting OptiPrep (60% aqueous iodixanol) with 0.25 M sucrose/10 mM Tris, pH 7.5. Aer an 18-hour centrifugation step at 100,000 x g, 12 density gradient fractions were collected, diluted in PBS and centrifuged at 100,000 x g for 3 hours. EVs from fractions 6 and 7 were combined and used as small EVs. To quantitate the size and concentration of EVs, nanoparticle tracking analysis was performed using a Particle Metrix ZetaView PMX 110. Cell numbers were gathered and viability checked at the end of conditioned media collection (48 h) to allow for calculation of EVs/cell/hour secretion rates.

### Mass spectrometry

20 µg of protein from vesicle preparations were first partially resolved by polyacrylamide gel electrophoresis approximately 1.5 cm using a 10% Novex precast gel. The region was excised and subjected to in-gel tryptic digestion to recover peptides. The resulting peptides were analyzed by a top-15 data dependent LC-MS/MS strategy using a QExactive plus mass spectrometer (ThermoFisher). Briefly, peptides were autosampled onto a 200 mm by 0.1 mm (Jupiter 3 micron, 300A), self-packed analytical column coupled directly to the mass spectrometer via a nanoelectrospray source and resolved using an aqueous to organic gradient with a 2-hour acquitision me. Both the intact masses (MS) and fragmentation paterns (MS/MS) of the peptides were collected utilizing dynamic exclusion to maximize depth of coverage. Resulting peptide MS/MS spectral data were searched using SEQUEST [35] against the canonical human protein database (Uniprot), to which common contaminants and reversed version of each protein had been added. Resulting peptide spectral matches (PSMs) were collated, filtered, and compared using Scaffold (Proteome Software).

### Proteomics Data Analysis

Differentially expressed proteins between KD and WT groups were determined by DESeq2[36]. Proteins were ranked by the significance test statistic from DESeq2 (Supplemental Table 1) and Gene Set Enrichment Analysis [37] was performed on the pre-ranked proteins against the Gene Ontology C5, Curated C2, and Hallmark gene sets from the Molecular Signatures Database (MSigDB v2022.1).

### STRING PPI analysis

The Search Tool for the Retrieval of Reciprocity Genes (STRING) database v11.5 [38] (http://string-db.org/) was used to predict the potential interactions between gene candidates at the protein level. The combined score of high confidence > 0.7 was used as the cut-off value in the STRING database. In PPI network figures, proteins showing no association with other proteins were removed.

### Western blot analysis

Protein concentrations of total cell and EV lysates were respectively determined utilizing Pierce BCA (Cat. 23225, Thermo Fisher) and Pierce Micro BCA (Cat. 23235, Thermo Fisher) Assays. For Western blots, 15 µg of TCLs or small EVs, were boiled in SDS-Page sample buffer for 5 min and loaded on 10% polyacrylamide gels. Proteins were transferred to nitrocellulose membranes for 1 h at 100 volts or 25 volts for overnight at 4°C. Membranes were blocked in 3% BSA diluted in Tris-buffered saline with 0.5% Tween 20 (TBST) for 4h at room temperature. Primary antibodies (from Abcam or Cell Signaling) were diluted in 3% BSA-TBST (SF3A3 (ab156873),1:5000; CD133/Prominin-1 (ab222782), 1:5000; EIF4A3 (ab180573), 1:1000; TSG101 (ab30871), 1:5000; and GAPDH (CST5174), 1:10000) and incubated overnight at 4C. Membranes were washed 3 times for 15 min in TBST and subsequently incubated with species-specific HRP-conjugated secondary antibodies (1:10000; Promega) in 3% BSA -TBST for 1h at room temp. All membranes were washed 3 times for 15 min in TBST and incubated with enhanced chemiluminescence (ECL) reagent (Thermo Scientific) for 1 min before being exposed to Amersham 680 imager (GE). Multiple exposures were taken for each blot to have the complete dynamic range for densitometry measurements, which were performed using the Analysis Gels feature of ImageJ (NIH).

## Acknowledgements

Funding was provided by NIH grant P01CA229123. We gratefully acknowledge feedback from the P01 group, the Weaver RNA group, and Adnan Shafiq. We thank Hayes McDonald in the Mass Spectrometry Research Center (MSRC) Proteomics Laboratory at Vanderbilt University for conducting the LC-MS/MS analysis.

## Figure and Table Legends

Table 1. Abbreviated list of GSEA pathways enriched in downregulated proteins in VAPA-KD small EVs. For each gene set, the pathway/function name, # of proteins, normalized enrichment score (NES), and false discovery rate (FDR) q value are listed.

Table 2. Abbreviated list of GSEA pathways enriched in upregulated proteins in VAP-A-KD SEVs. For each gene set, the pathway/function name, # of proteins, normalized enrichment score (NES), and false discovery rate (FDR) q value are listed.

**Supplemental Figure 1:**
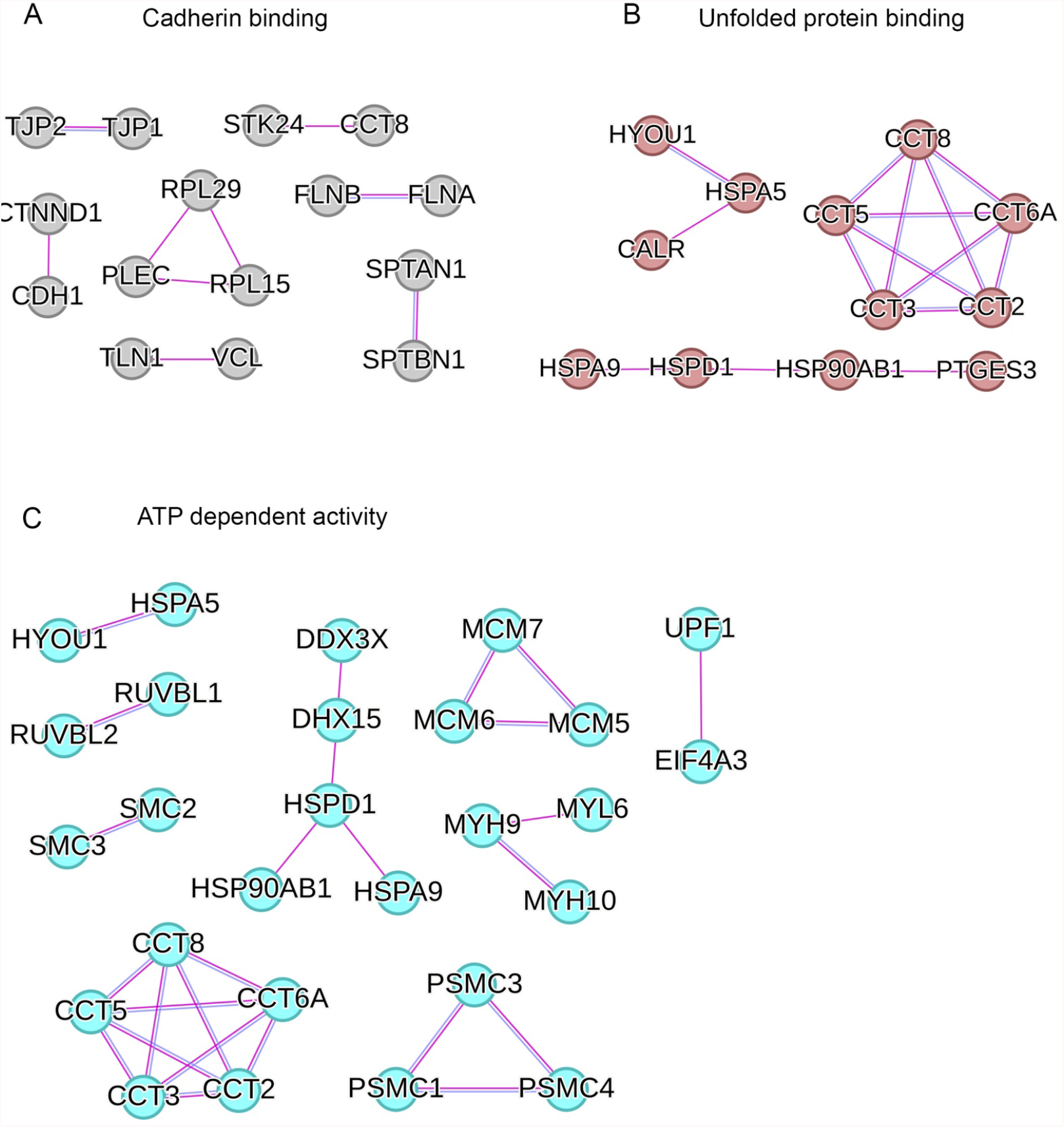
Additional PPI networks with downregulated proteins in VAP-A-KD SEVs. A. PPI networks by STRING for downregulated proteins in GOMF_Cadherin binding, GO:0045296. B. PPI networks by STRING for downregulated proteins in GOMF_Protein folding chaperone, GO:0044183. C. PPI networks by STRING for downregulated proteins in GOMF_ATP dependent activity, GO:0140657

**Supplemental Figure 2:**
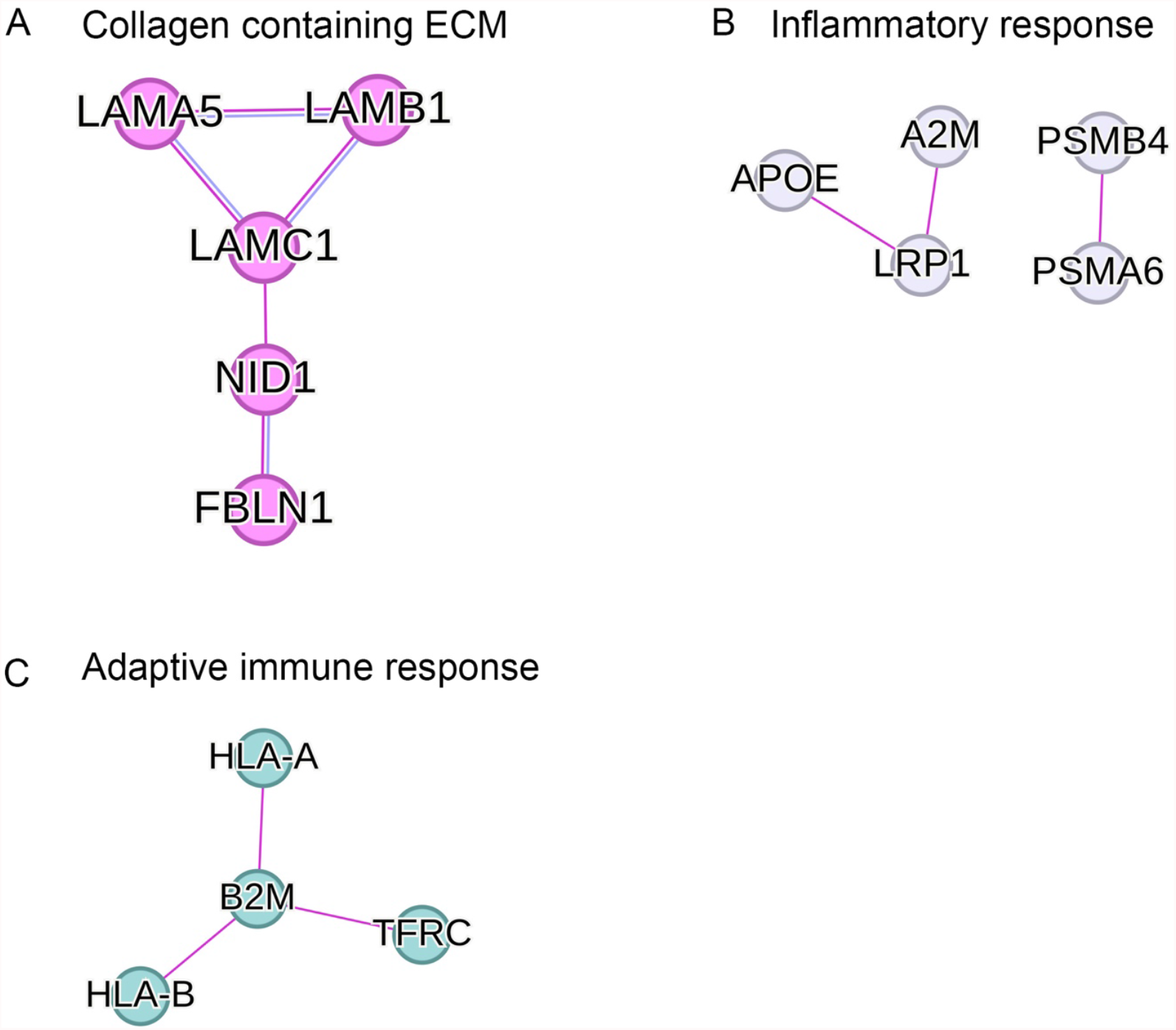
PPI networks for select upregulated proteins in VAP-A-KD SEVs. A. PPI networks by STRING for upregulated proteins in Collagen containing ECM proteins. B. PPI networks by STRING for upregulated proteins in GSEA category for Collagen containing inflammatory response. C. PPI networks by STRING for upregulated proteins in GSEA category for Collagen containing adapve immune response.

Supplemental table 1: List of all proteins with their spectral counts and analyzed data found in control (Sc) and VAP-A-KD (KD1 and KD2) SEVs.

Supplemental table 2: List of all downregulated proteins found in GSEA gene sets in VAP-A-KD SEVs.

Supplemental table 3: List of all upregulated proteins found in GSEA gene sets in VAP-A-KD SEVs.

## Associated data

The mass spectrometry proteomics data have been deposited to the ProteomeXchange Consortium via the PRIDE partner repository with the dataset identifier PXD043796.

